# Fatty acid binding proteins shape the cellular response to activation of the glucocorticoid receptor

**DOI:** 10.1101/2021.07.02.450968

**Authors:** Bonan Liu, Indu R. Chandrashekaran, Olga Ilyichova, Damien Valour, Fabien Melchiore, Chantal Bourrier, Adeline Giganti, Jean-Philippe Stephan, Catherine Dacquet, Patrick Genissel, Willy Gosgnach, Richard J. Weaver, Christopher J.H. Porter, Martin J. Scanlon, Michelle L. Halls

**Affiliations:** Drug Discovery Biology, Monash Institute of Pharmaceutical Sciences, Monash University, 381 Royal Parade Parkville, Victoria 3052, Australia; Drug Delivery, Disposition and Dynamics, Monash Institute of Pharmaceutical Sciences, Monash University, 381 Royal Parade Parkville, Victoria 3052, Australia; Medicinal Chemistry, Monash Institute of Pharmaceutical Sciences, Monash University, 381 Royal Parade Parkville, Victoria 3052, Australia; ARC Training Centre for Fragment-Based Design, Monash Institute of Pharmaceutical Sciences, Monash University, 381 Royal Parade Parkville, Victoria 3052, Australia; Institut de Recherches Servier, 78290 Croissy-Sur-Seine, 50, rue Carnot, 922284 Cedex, France; Institut de Recherches Servier, 92210 Suresnes, 50, rue Carnot, 922284 Cedex, France; Institut de Recherches Internationales Servier, 50, rue Carnot, 922284 Cedex, France

**Keywords:** Fatty acid binding protein, glucocorticoid receptor, nuclear hormone receptor, drug discovery, transcription

## Abstract

Glucocorticoids are steroid hormones that are essential for life in mammals. Therapeutically, they are some of the most cost-effective drugs for the treatment of inflammatory diseases ranging from skin rashes to COVID-19, but their use is limited by adverse effects. Glucocorticoids exert their effects via the glucocorticoid receptor, a type I nuclear hormone receptor which modulates gene expression. The transcriptional activity of some related, but nuclear restricted, type II nuclear hormone receptors can be enhanced by a family of intracellular transport proteins, the fatty acid binding proteins (FABPs). We find that the transcriptional activity of the GR can be altered by a sub-set of FABP family members dependent on the GR-ligand. The ability of some FABPs to selectively promote or limit the transcriptional activity of the GR in a ligand-dependent manner could facilitate the discovery of drugs that narrow GR activity to only the desired subset of therapeutically relevant genes.

## Introduction

Glucocorticoids are steroid hormones that are essential for life in mammals. They play a key role in physiological processes ranging from immune response to metabolism, cardiovascular function and development. Therapeutically, both endogenous and synthetic glucocorticoids are some of the most used and cost-effective drugs for the treatment of inflammatory diseases such as asthma, skin rashes, rheumatoid arthritis and COVID-19^1, 2^. However, glucocorticoid therapy is limited by frequent and severe adverse effects including osteoporosis, fat redistribution, hyperglycaemia, cardiovascular disease and infections^3^. For example, patients on long-term oral glucocorticoid therapy have an adjusted odds ratio for cardiovascular events of 2.56^4^, and 30-50% of this population will experience bone fractures^5^. The challenge remains to fully appreciate mechanisms of drug action and ADME (absorption, distribution, metabolism and excretion) governing the desired pharmacological effects in addition to the off-target effects that currently limit their use.

Glucocorticoids are synthesised from a cholesterol precursor, and freely diffuse across the plasma membrane due to their lipophilic nature. In the cytosol, glucocorticoids bind their target nuclear hormone receptor (NHR), the glucocorticoid receptor (GR). The GR exists within a multi-protein complex in the cytosol; binding of glucocorticoids induces a conformational change in the GR and a rearrangement of the GR-protein complex, which ultimately facilitates translocation of the activated GR to the nucleus. In the nucleus, the GR can affect approximately 10% of all genes through direct DNA binding (i.e. by acting as a transcription factor), or by modulating the function of other transcription factors^6^. GR-DNA binding is terminated by exportins and calreticulin, which regulate the nuclear export of the GR back to the cytoplasm^7^. It is the balance between nuclear import and export which determines the proportion of GR in the nucleus (compared to the cytosol) and therefore the strength of the transcriptional activity of the GR^7, 8^. The transcriptional output of the activated GR is influenced by physical interactions with other proteins recruited to the receptor following ligand binding and nuclear import^9^. The composition of the GR-protein complex is dependent on the cell-specific expression of different proteins and the conformational changes in the GR induced by ligand binding, DNA binding and post-translational modification^10^. The activity of the GR and which genes are affected therefore changes in a cell and context specific manner.

We and others have previously shown that a class of intracellular lipid binding proteins, the fatty acid binding proteins (FABPs), can increase the transcriptional activity of another family of NHRs, the peroxisome proliferator-activated receptors (PPARs)^11, 12, 13, 14, 15, 16^. This occurs in a ligand- and PPAR-selective manner and is associated with an increase in the nuclear localisation of the FABP^11, 13, 15^. The mechanism by which FABPs promote activation of PPAR appears to be driven by a conformational change in the FABP upon FABP-ligand binding that modulates the electrostatic surface of the FABP and promotes activation of the PPAR transcriptional complex^12, 13, 15, 17^. Given their role in transporting poorly water-soluble ligands throughout the cell, and their ability to modulate the activity of NHRs, we hypothesised that FABPs may also modulate the transcriptional activity of the GR. FABPs have very specific cell and tissue expression patterns in healthy individuals, and this varies with the onset of disease^18, 19^. As such, any influence of FABPs on GR activity could provide significant insight into the role of intracellular trafficking proteins in determining pharmacokinetic/pharmacodynamic relationships and drug action at NHRs. Here, we probed the potential effect of five FABPs on the transcriptional activity of the GR: FABP1 (liver, L-FABP), FABP2 (intestine, I-FABP), FABP3 (heart, H-FABP), FABP4 (adipocyte, A-FABP) and FABP5 (brain, B-FABP or epidermal, E-FABP). We find that the transcriptional activity of the GR is enhanced in the presence of FABP1 or FABP4, in a ligand-dependent manner. The FABP-dependent increase in GR transcriptional activity is associated with increased nuclear localisation and increased proximity of both the GR and FABP, due to slowed nuclear egress and the dexamethasone-induced increase in GR transcriptional activity is limited to genes containing a glucocorticoid response element (GRE) site. However, when examined at the level of the transcriptome, we surprisingly find that FABP4 co-expression reverses dexamethasone-induced changes in gene expression. These data suggest that expression patterns of FABPs have an important role in defining the cell response to activation of the GR.

## Results

### FABPs can potentiate the transcriptional activity of GR but not MR

The endogenous glucocorticoids, cortisol (the predominant form in humans) and corticosterone, can bind and activate both the GR and the mineralocorticoid receptor (MR). In addition to a form of cortisol used therapeutically (hydrocortisone), various synthetic glucocorticoids have been developed as therapeutics e.g. dexamethasone and prednisolone. The synthetic glucocorticoids are based on the structure of the endogenous glucocorticoids, cortisol and hydrocortisone, but with modifications to optimise their pharmacokinetics, bioavailability, and specificity for the GR over the MR.

We initially tested whether expression of FABPs 1-5 could affect the transcriptional activity of the GR in transfected COS-7 cells following stimulation with hydrocortisone, dexamethasone or prednisolone. COS-7 cells were used as they do not endogenously express detectable levels of FABPs 1-5 (Fig S1). The simplest form of GR-DNA interaction is the binding of the GR to genomic glucocorticoid binding sites containing a glucocorticoid response element (GRE); a 15 base pair sequence motif of two imperfect inverted palindromic repeats of six base pairs, separated by a three base pair spacer^1^. We therefore assessed the level of GR transactivation using a GRE reporter gene assay.

Hydrocortisone caused a concentration-dependent increase in GR transactivation in COS7 cells expressing the GR, but we observed no change in this activity with co-transfection of FABP1-5 (Fig 1A). In contrast, the maximal GR transactivation following stimulation of cells with dexamethasone was increased with cotransfection of FABP4, but not FABP1, FABP2, FABP3, or FABP5 (Fig 1B). FABP1 or FABP4 transfection also increased the maximal GR transactivation in response to prednisolone, compared to a vector control (Fig 1C). To confirm these responses were due to activation of the transfected GR and not endogenous MR in COS-7 cells, we performed the same experiment using EC_80_ concentrations of ligand in COS-7 cells transfected with the MR and one of FABP1-5. In contrast to GR transfection, there was no change in the activation of GRE-driven transcription in COS-7 cells co-transfected with MR and any of the FABPs in response to hydrocortisone, dexamethasone or the MR endogenous ligand, aldosterone (Fig S2). COS-7 cells endogenously express a small amount of GR (Fig 1D); transfection with FABP4 alone increased the maximal GR transactivation in response to dexamethasone and prednisolone, but this was further increased with GR co-transfection (Fig 1E-F). Taken together, this suggests that the dexamethasone-induced increase in GRE transcription is dependent on the GR.

**Figure 1.**
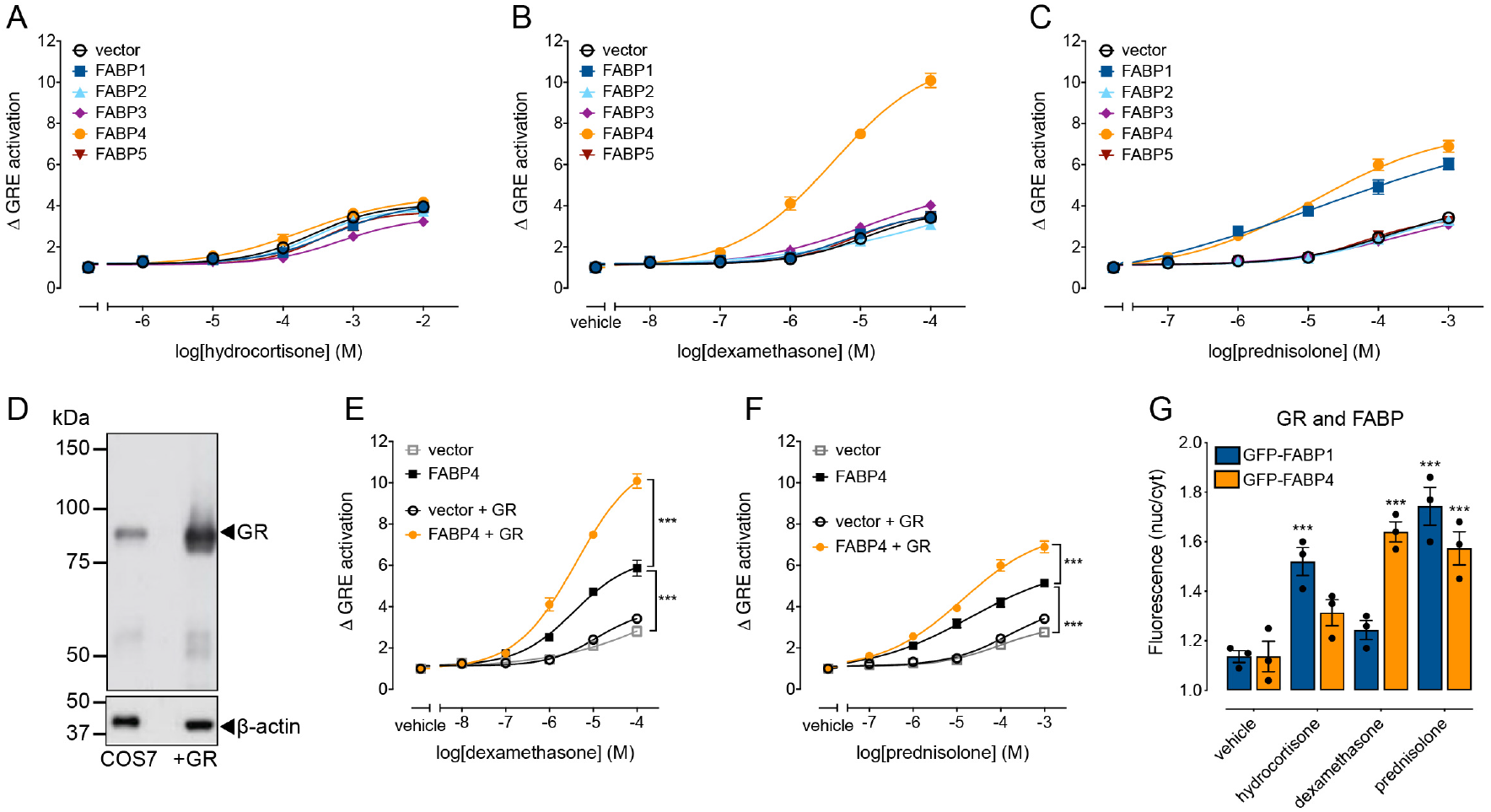
Co-transfection of FABP1 or FABP4 increases GR-dependent transcriptional activity in a ligand-dependent manner. GRE-transcriptional activity of GR in the absence or presence of FABP1, FABP2, FABP3, FABP4 or FABP5 in response to vehicle control or increasing concentrations of (A) hydrocortisone (B) dexamethasone or (C) prednisolone, was assessed using a reporter gene in COS-7 cells after 24 h treatment (n=3). (D) Expression of the GR in naïve COS-7 cells or following transfection with GR determined by immunoblotting. Transcriptional activity of the endogenous or transfected GR in the presence of FABP4 in response to (E) dexamethasone or (F) prednisolone (n=3). (G) FABP translocation to the nucleus in COS-7 cells transfected with GR and GFP-FABP1 or GFP-FABP4 following 24 h treatment with vehicle or an EC_80_ concentration of hydrocortisone, dexamethasone or prednisolone (n=3). Data are mean ± SEM from *n* independent experiments, as stated. For concentration-response curves, symbols show means and error bars, S.E.M. *** p<0.001, two-way ANOVA with Sidak’s multiple comparison test. For bar graphs, bars show the mean, error bars the S.E.M. and symbols show the independent data points for each experiment. *** p<0.001, two-way ANOVA with Dunnett’s multiple comparisons test.

We have previously shown that FABP-induced increases in PPAR transcriptional activity are associated with an accumulation of FABP in the nucleus^11, 12^. We therefore measured the ability of an EC_80_ concentration of hydrocortisone, dexamethasone or prednisolone to affect the cellular distribution of GFP-tagged FABP (Fig 1G). We quantified the cytosolic vs nuclear distribution of GFP-FABP1 or GFP-FABP4, as transfection of cells with these FABPs (but not FABP2, FABP3 or FABP5) enhanced the GRE-transcriptional activity of the GR in a ligand-dependent manner. Both dexamethasone and prednisolone, but not hydrocortisone, caused a redistribution of GFP-tagged FABP4 to the nucleus compared to treatment with vehicle control. This was consistent with the relative ability of these ligands to affect GRE-transcriptional activity following FABP4 transfection (i.e. no effect on responses to hydrocortisone, but enhancement of dexamethasone and prednisolone activity; Fig 1A-C). This is consistent with a requirement of prolonged nuclear localisation of FABP for enhanced PPAR activity^11, 13, 15^. In contrast, both hydrocortisone and prednisolone, but not dexamethasone, caused a redistribution of GFP-tagged FABP1 to the nucleus compared to treatment with vehicle control. While this result was consistent with the relative ability of dexamethasone and prednisolone to affect GRE-transcriptional activity following FABP1 transfection, it was not the case for hydrocortisone (Fig 1A-C). We have previously reported similar observations for the PPAR system, whereby some ligands can promote FABP redistribution to the nucleus, without any detectable effect on reporter gene activity^11^.

### FABPs only enhance GRE-transcriptional activity in response to selected glucocorticoids in COS-7 cells

Given the role of synthetic glucocorticoids in the treatment of a wide range of pathologies, we then assessed whether FABP1 or FABP4 could affect the GRE-transcriptional activity of a wider range of ligands (Fig 2). There was no effect of FABP1 or FABP4 co-transfection on the GRE-transcriptional activity in response to increasing concentrations of the endogenous glucocorticoid, cortisone (Fig 2A). In contrast, the GRE-transcriptional activity in response to the synthetic glucocorticoids, β-methasone, budesonide and methylprednisolone, was affected in the presence of FABPs. While only FABP4 enhanced the GRE-transcriptional activity of β-methasone (Fig 2B) and budesonide (Fig 2C), both FABP1 and FABP4 increased the transcriptional activity of methylprednisolone (Fig 2D). Interestingly, there was also no effect of FABP co-transfection on the GRE-transcriptional activity induced by prednisone (Fig 2E).

**Figure 2.**
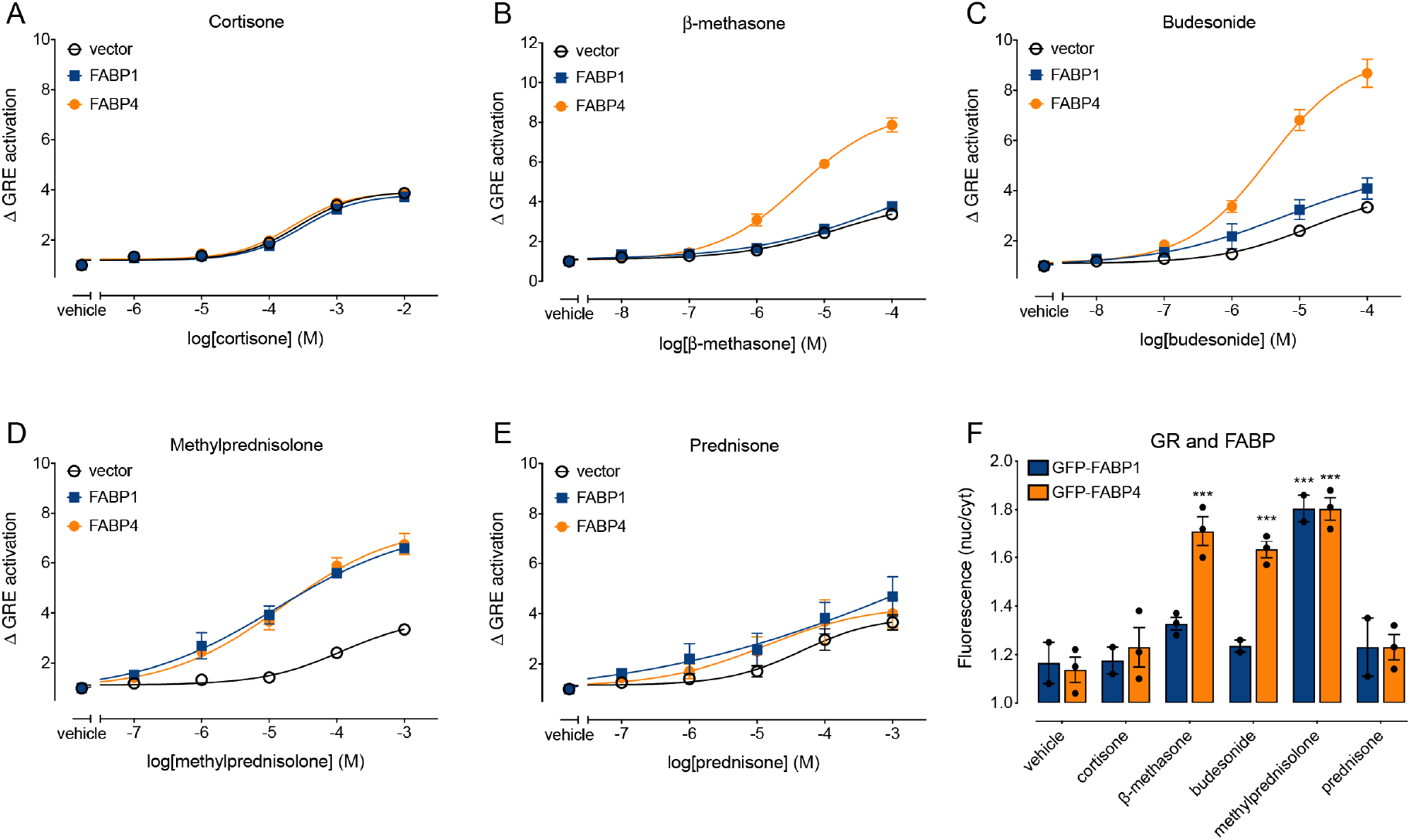
FABP1 and FABP4 affect the activation of GR in response to selected glucocorticoids. Transcriptional activity of GR in the absence or presence of FABP1 or FABP4 was assessed using a GRE reporter gene in COS-7 cells following 24 h treatment with vehicle or increasing concentrations of (A) cortisone, (B) β-methasone, (C) budesonide, (D) methylprednisolone or (E) prednisone (n=3). Symbols are means and error bars SEM. (F) FABP translocation to the nucleus in COS-7 cells transfected with GR and GFP-FABP1 or GFP-FABP4 following 24 h treatment with vehicle or an EC_80_ concentration of cortisone, β-methasone, budesonide, methylprednisolone or prednisone (n=3). Bars are means, error bars SEM and symbols show individual data points for each experiment. *** p<0.001 versus vehicle control, two-way ANOVA with Dunnett’s multiple comparison test.

We next measured the ability of an EC_80_ concentration of each of the GR ligands to affect the cellular distribution of a GFP-tagged FABP (Fig 2F). Only β-methasone, budesonide and methylprednisolone caused a redistribution of GFP-tagged FABP to the nucleus. This occurred in a ligand-specific and FABP-specific manner and was consistent with the ability of the ligand/FABP pair to increase GRE-transcriptional activity. Specifically, β-methasone and budesonide only increased nuclear localisation of GFP-FABP4 (but not GFP-FABP1), whereas methylprednisolone increased the nuclear localisation of both GFP-FABP1 and GFP-FABP4 (Fig 2F).

In addition to regulating the transcriptional activity of PPARs, FABPs have been traditionally recognised as “ligand transporters” as they are required to shuttle water-insoluble fatty-acids from the plasma membrane to their receptor in the nucleus. FABPs can therefore directly bind fatty acid ligands; for example, FABP1 binds the endogenous fatty acid, oleic acid (oleate) with a K_D_ of 0.58 μM and the synthetic PPARa agonist GW7647 with a K_D_ of 0.12 μM, as determined by isothermal titration calorimetry (ITC)^12^. We therefore assessed whether FABPs could also bind GR ligands using a fluorescence polarisation assay (Table 1). Although we could observe binding of some GR ligands to some FABPs, this occurred only at very high concentrations of the GR agonists. For the two FABPs that increased transcriptional activity of the GR, FABP1 and FABP4, calculated pKi values for the GR ligands were in the high micromolar to millimolar range (Table 1). In contrast, we find that dexamethasone, cortisone and methylprednisolone have affinities (Ki) of 4.8 nM, 1.5 μM and 2.7 nM, respectively, for the GR (Table S1). Therefore, FABPs are highly unlikely to directly bind GR ligands *in vivo*.

**Table 1.** Binding affinity of GR ligands (endogenous and synthetic) for FABPs. Competition binding between GR ligands and BODIPY-C4,C9 fatty acid (200 nM) for purified FABP1, FABP2, FABP3, FABP4 or FABP5. The table shows pKi values as means ± standard error of the mean from three experiments performed in triplicate. N.D., not determined due to ambiguous fit of the curve.

| Ligand | FABP1 | FABP2 | FABP3 | FABP4 | FABP5 |
| --- | --- | --- | --- | --- | --- |
| Dexamethasone | 4.56±0.03 | N.D. | 5.25±0.01 | 4.13±0.09 | 4.59±0.08 |
| Hydrocortisone | 4.41±0.02 | N.D. | 5.11±0.03 | 3.92±0.13 | 4.66±0.16 |
| Prednisolone | 4.32±0.14 | 4.96±0.65 | 4.15±0.34 | 2.94±0.33 | 4.28±0.46 |
| Cortisone | 3.52±0.08 | 4.78±0.35 | 3.97±0.10 | 3.01±0.13 | 3.89±0.34 |
| β-methasone | 4.04±0.05 | 4.14±0.08 | N.D. | N.D. | 6.52±0.63 |
| Budesonide | 4.07±0.14 | N.D. | 7.24±0.11 | N.D. | 4.72±0.92 |
| Methylprednisolone | 4.10±0.13 | 5.47±0.87 | 3.93±0.08 | 2.70±0.42 | 3.62±0.11 |
| Prednisone | 3.54±0.10 | N.D. | 5.12±0.02 | 4.45±0.22 | 5.03±0.14 |

### An FABP-induced increase in GRE activity correlates with increased GR-FABP FRET in the nucleus

We have previously shown that FABP-induced increases in PPAR transcriptional activity depend on a close and prolonged association between the FABP and PPAR in the nucleus^11, 12^. PPARs are localised to the nucleus, and the prolonged association between PPAR and FABP requires translocation of the FABP to the nucleus. In contrast, the GR is found distributed between the nucleus and the cytosol but moves to the nucleus following activation. In order to determine where the GR/FABP interaction first occurs, we measured FRET between GR-dsRed and GFP-tagged FABP1 or FABP4 in the nucleus compared to the cytosol of cells in response to EC_80_ concentrations of hydrocortisone, dexamethasone or prednisolone (Fig 3). GR-dsRed caused a similar increase in GRE-transcriptional activity in response to dexamethasone, compared to the untagged GR (Fig S3).

**Figure 3.**
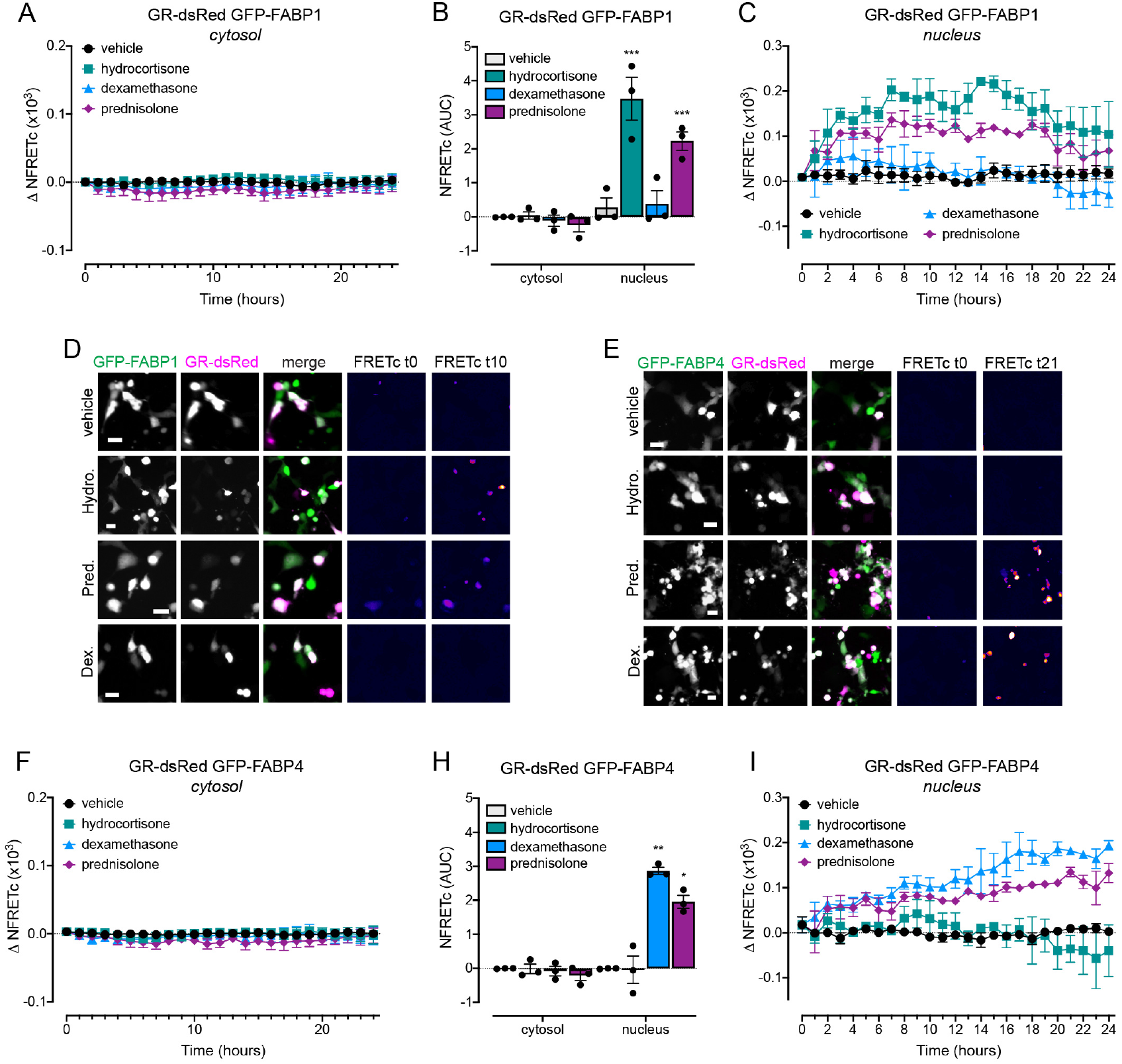
Ligand-induced FRET between GR-dsRed and FABP-GFP occurs in the nucleus. Time course of FRET between GR-dsRed and GFP-FABP1 (A-D) or GFP-FABP4 (E-I) in COS-7 cells in response to vehicle or an EC_80_ concentration of hydrocortisone, dexamethasone or prednisolone (n=3). A and F show the ligand-induced change in normalised corrected FRET (NFRETc) in the cytosol (image region automatically defined by subtracting the nuclear region from the whole cell region). Symbols are means and error bars SEM. B and H show the area under the curve (AUC) calculated from the time course data in A and C or F and I, respectively. Bars are means, error bars are S.E.M., and symbols show individual data points from each experiment. * p<0.05, ** p<0.01 and *** p<0.001 versus vehicle control, two-way ANOVA with Dunnett’s multiple comparisons test. C and I show the ligand-induced change in normalised corrected FRET (NFRETc) in the nucleus (region defined by a nuclear Hoescht stain). Symbols are means and error bars SEM. D and E show representative GFP, dsRed, merge and pseudocoloured FRET images (calculated using the FRET Analyzer plugin for the Fiji distribution of Image J). Pseudocolour FRET images are shown at baseline (t0) and when FRET signal plateaus (at 10 hours after drug addition for FABP1 or at 21 hours after drug addition for FABP4). Dark blue shows low corrected FRET (FRETc) and yellow/white shows high FRETc; scale bar is 20 μm; Hydro., hydrocortisone, Pred., prednisolone and Dex., dexamethasone.

There was no change in FRET between GR-dsRed and GFP-FABP1 in the cytosol in response to any ligand tested (Fig 3A,B). Both hydrocortisone and prednisolone, but not dexamethasone, caused an increase in nuclear FRET between GR-dsRed and GFP-FABP1 (Fig 3B-D). The ability of a particular ligand/FABP pair to increase FRET between GR-dsRed and GFP-FABP1 was consistent with whether the ligand caused a redistribution of FABP1 to the nucleus (Fig 1G). Similarly, there was no change in FRET between GR-dsRed and GFP-FABP4 in the cytosol in response to any ligand tested (Fig 3E-G). Dexamethasone and prednisolone, but not hydrocortisone, caused an increase in FRET in the nucleus between GR-dsRed and GFP-FABP4 (Fig 3E,G-I). This was consistent with the redistribution of FABP4 to the nucleus and the increase in the GRE-transcriptional activity in the presence of FABP4 in response to dexamethasone and prednisolone, but not hydrocortisone (Fig 1A-C,G).

Increased GR GRE-transcriptional activity in the presence of FABPs therefore occurs in a ligand dependent manner. It is associated with an increased proximity between the FABP and GR which is restricted to the nucleus, suggesting that the interaction occurs after GR activation and translocation.

### Prolonged nuclear residence of GR and FABP4 is due to decreased nuclear egress

We then sought to determine the mechanism underlying the increased accumulation of both GR and FABP in the nucleus using FABP4 as a model, as this FABP caused the largest potentiation of dexamethasone-induced activity. For two proteins that are usually co-distributed between the cytosol and nucleus, an accumulation in the nuclear region could be due to either an increase in nuclear entry, or a decrease in nuclear egress. To differentiate between these two processes (nuclear entry vs egress) we used fluorescent recovery after photobleaching (FRAP) (Fig 4).

**Figure 4.**
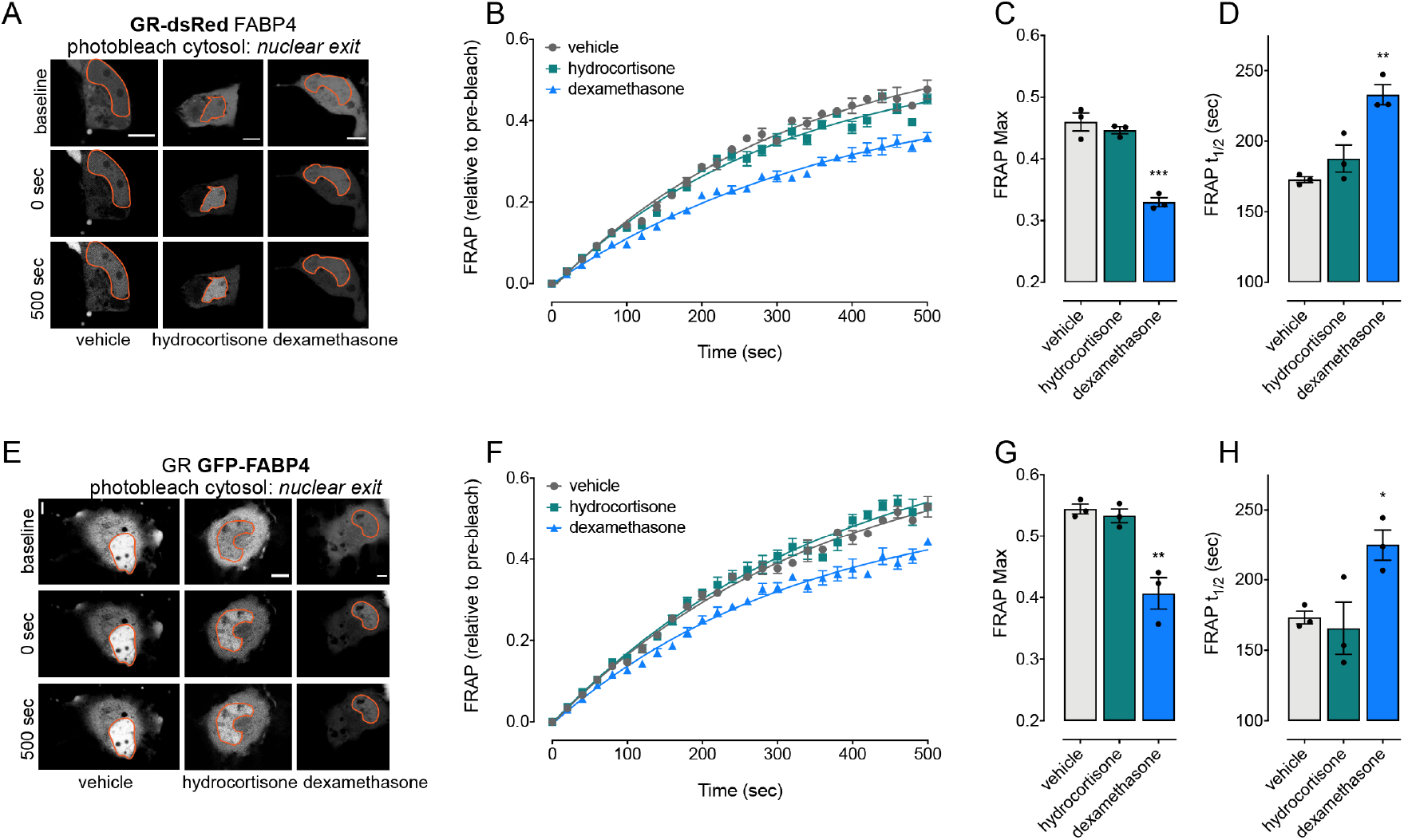
Enhanced GR transcriptional activity by FABP4 is associated with decreased nuclear egress. COS-7 cells were treated with vehicle, hydrocortisone or dexamethasone for 6 h, before photobleaching of the cytosol. Fluorescence recovery was monitored every 20 sec. (A-D) GR mobility was monitored in cells co-transfected with GR-dsRed and FABP4 (n=3). (E-H) FABP4 mobility was monitored in cells co-transfected with GR and FABP4-GFP (n=3). A and E show representative images with the nucleus outlined in orange. Scale bar is 10 μm. B and F show FRAP time course, symbols are means and error bars SEM. C and D show maximum recovery. G and H show the half time of recovery. Bars are means, error bars SEM and symbols show individual data points for each experiment. ** p<0.01 versus vehicle control, one-way ANOVA with Sidak’s multiple comparison test.

To monitor changes in the rate of nuclear entry, we photobleached the nucleus, and then monitored the rate of recovery of either GR-dsRed or GFP-FABP4 fluorescence back into the nucleus. We observed no change in the maximal fluorescence recovery, or in the rate of fluorescence recovery in the nucleus for either GR-dsRed or GFP-FABP4 in response to dexamethasone (activating) or hydrocortisone (control) (Fig S4). To monitor changes in the rate of nuclear egress, we photobleached the cytosol, and then monitored the rate of recovery of either GR-dsRed or GFP-FABP4 fluorescence back into the cytosol (Fig 4). Dexamethasone stimulation caused a smaller fluorescence recovery and an increased half-time of fluorescence recovery into the cytosol in cells co-transfected with either GR-dsRed (with untagged FABP4, Fig 4A-D) or GFP-FABP4 (with untagged GR) (Fig 4E-H). There was no difference in fluorescence recovery in these same cells treated with hydrocortisone compared to the vehicle control.

Therefore, the increase in nuclear accumulation and association between GR and FABP4 is controlled by a slower rate of nuclear egress, and not an increase in nuclear entry. This occurs in a ligand-specific manner, and is consistent with a lack of FRET between the two proteins in the cytosol (Fig 3).

### FABP4 co-transfection enhances the dexamethasone-induced expression of a sub-set of genes involved in GR signalling

To examine the consequences of increased GRE-mediated gene transcription, we used a commercial qPCR array specific to GR signalling (Fig 5). It includes the GR and co-transcription factors, in addition to GR target genes identified by chromatin immunoprecipitation (ChIP). Therefore, the array principally includes target genes with GRE sites.

**Figure 5.**
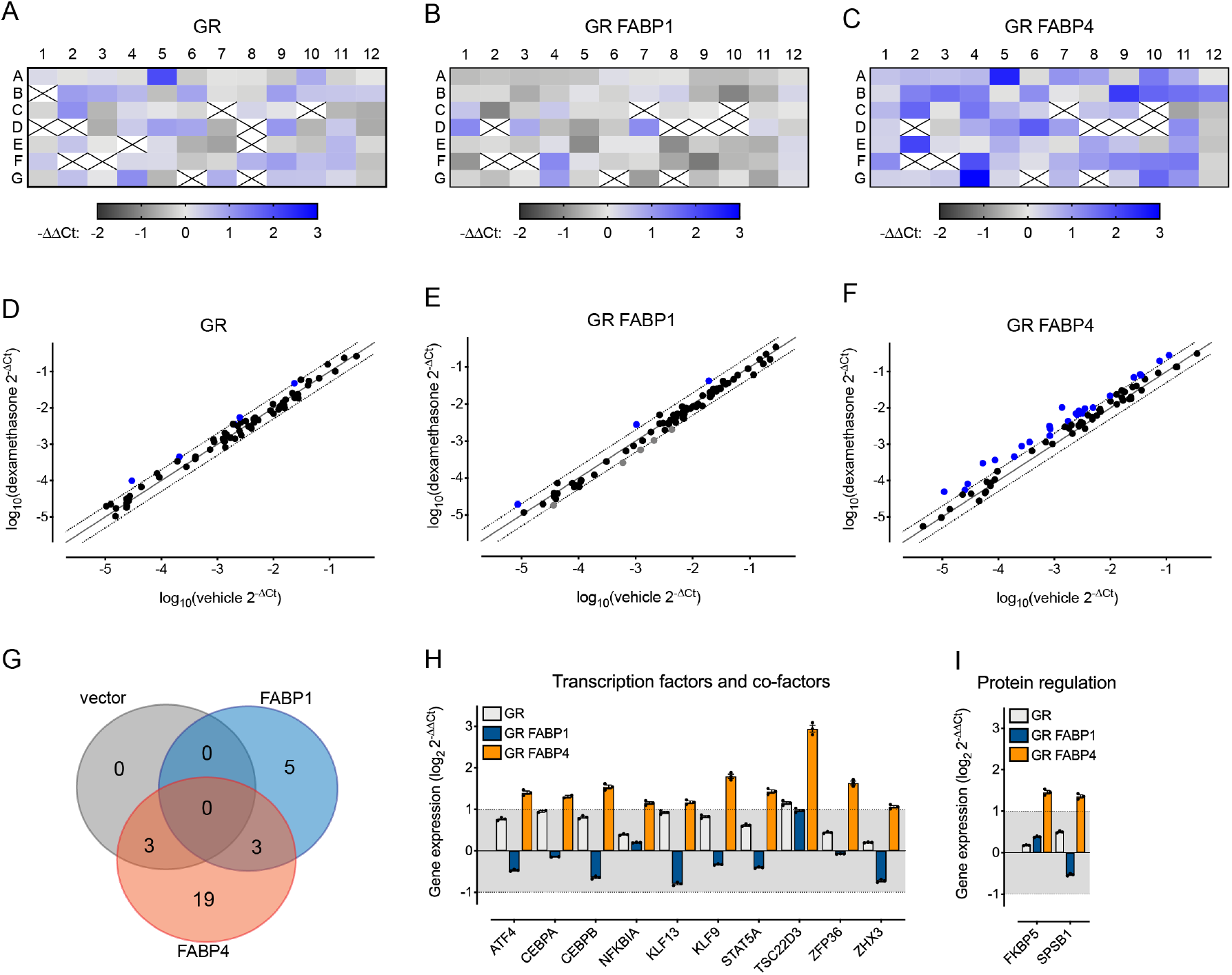
FABP4 increases GR-mediated gene transcription in response to dexamethasone. Heat maps of the dexamethasone-induced change in expression of 83 GR-associated genes after 24 h in COS-7 cells co-transfected with GR and (A) vector control, (B) FABP1 or (C) FABP4 (n=3). Scatter plots showing the change in gene expression in response to vehicle or dexamethasone in COS-7 cells co-transfected with GR and (D) vector control, (E) FABP1 or (F) FABP4; genes with at least a 2-fold change are represented as coloured dots (blue shows increase, grey shows decrease; black shows unchanged). Dots show average of n=3. (G) Venn diagram showing the number of genes that changed at least 2-fold in response to dexamethasone for each transfection condition. (H) Genes that encode transcription factors or co-factors with an average 2-fold change in expression in cells transfected with GR and FABP4 in at least 2 experiments. (I) Genes that affect protein regulation that had an average 2-fold change in expression in cells transfected with GR and FABP4 in at least 2 experiments. Bars are means, error bars SEM and symbols show individual data points for each experiment. Shaded area (between −1 and 1) indicates genes with less than a 2-fold change in response to dexamethasone.

COS-7 cells were co-transfected with GR and either pcDNA (control), FABP1 (negative control) or FABP4, and then treated with vehicle or dexamethasone for 24 h. We calculated the log2 fold change in Ct value in response to dexamethasone relative to vehicle and the housekeeping genes HPRT1 and RPLP0 (Table S1). The averaged data from three biological replicates is shown as a heat map for GR co-transfected with pcDNA (Fig 5A), FABP1 (Fig 5B) or FABP4 (Fig 5C). Over the entire dataset, we observed at least a 2-fold change in response to dexamethasone compared to the vehicle control in 36 out of a total of 84 genes. No signal was observed for 9 genes, and 39 genes showed changes in response to dexamethasone which did not reach our significance threshold (Table S1).

COS-7 cells co-transfected with GR and FABP4 showed a larger number of genes that were upregulated in response to dexamethasone (25 genes up, 0 genes down), compared to cells co-transfected with vector (3 genes up, 0 genes down) or FABP1 (3 genes up, 5 genes down) (Fig 5D-F). Moreover, of these genes that were differentially regulated in response to dexamethasone, 19 genes were uniquely regulated in cells co-transfected with FABP4, compared to vector control (0 unique genes) or FABP1 (5 unique genes) (Fig 5G). This is consistent with our GRE reporter gene assay showing an increase in GRE-transcriptional activity in response to dexamethasone in cells co-transfected with FABP4, compared to GR alone or cells co-transfected with FABP1 (Fig 1).

Of the genes differentially regulated by dexamethasone in cells co-transfected with FABP4, approximately half were identified as transcription factors or co-factors (Fig 5H) or involved in protein folding and regulation (Fig 5I). The remaining genes encode receptor ligands, transporters and proteins involved in cellular trafficking, signalling, the stress response, and regulation of collagen (Fig S5).

### FABP4 co-transfection narrows the transcriptional focus of the GR

The GR can regulate gene transcription by multiple mechanisms. The simplest form of GR-DNA binding involves the binding of GR homodimers to consensus GRE sites, as examined in Figure 5. However, the GR can also act as a monomer and bind to half-GRE sites, repress gene transcription by binding to an inverted repeat GR-binding site, or interact with other transcription factors^1^. To determine how FABPs could influence GR-mediated gene transcription as a whole, we performed RNA-Seq.

COS-7 cells were co-transfected with GR and either pcDNA (control), FABP1 (negative control) or FABP4, and then treated with vehicle or dexamethasone for 24 h. RNA was isolated, and the transcriptome determined by RNA-Seq. Of the genes identified, 18,385 were known genes and 4,286 were novel genes; 15,882 known genes could be mapped to gene IDs. The fold change in gene expression in response to dexamethasone (compared to vehicle control) was determined for each gene, and the entire gene list input into Ingenuity Pathway Analysis (IPA) software. After removing genes where there was little change in response to dexamethasone (removed log_2_(RPKM) values between −0.58 and 0.58) or genes that were identified at very low expression levels (removed RPKM intensity values between −0.1 and 0.1), 1,868 genes remained from cells co-transfected with GR and vector, 1,913 for GR and FABP1, and 772 genes for GR and FABP4.

To validate our data, we used IPA to confirm that the most likely upstream regulator of this set of genes was dexamethasone. The network of genes that was up-regulated and down-regulated in the control transfection condition (GR with vector) in response to dexamethasone is shown in Fig 6A. The GR-mediated transcriptional profile is highly cell type specific, highlighted by limited overlap in GR binding sites in different cell types and tissues^20, 21, 22, 23^. Despite this, of the genes included in the dexamethasone-regulated network we found that 23 out of 25 of the genes that were expected to be up-regulated, *were* upregulated by dexamethasone in our dataset (Fig 6B, S6A-B); similarly, 6 out of the 11 genes expected to be down-regulated, were down-regulated by dexamethasone in our dataset (Fig S6C-D).

**Figure 6.**
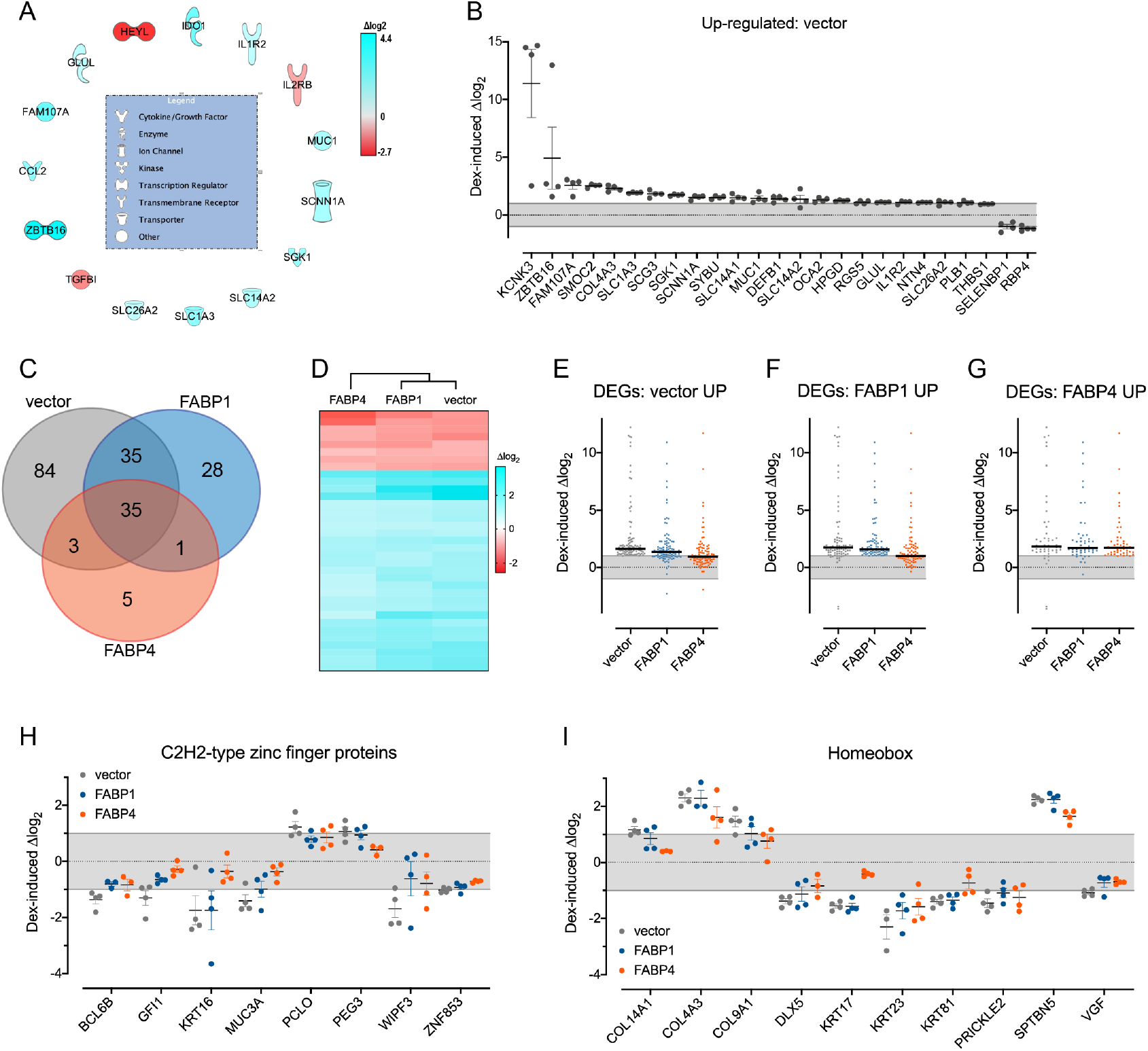
FABP co-transfection changes the transcriptomic profile of GR activated by dexamethasone. COS-7 cells co-transfected with GR and vector (control), FABP1 or FABP4 were treated with vehicle or dexamethasone for 24 h, prior to RNA isolation, sequencing and transcriptomic analysis (n=4). (A) Heatmap of the dexamethasone-signature genes (IPA network) that were differentially regulated by dexamethasone in the GR/vector transcriptome. (B) Dexamethasone-induced change in expression of genes in COS-7 cells co-transfected with GR/vector, that were expected to be upregulated by dexamethasone. Symbols show the data points from each biological replicate, bars show mean and SEM, grey shading indicates less than a 2-fold change in response to dexamethasone. (C) Venn diagram showing the number of genes that had at least a 2-fold change (differentially expressed genes) in response to dexamethasone in cells co-transfected with GR/vector, GR/FABP1 or GR/FABP4. (D) Hierarchical clustering of differentially expressed genes following dexamethasone treatment of cells co-transfected with GR/vector, GR/FABP1 or GR/FABP4. (E-G) The average change in differentially-expressed genes (DEGs) in response to dexamethasone across all transfection conditions, with a focus on genes that were up-regulated more than 2-fold in (E) GR/vector cells, (F) GR/FABP1 cells and (G) GR/FABP4 cells. Symbols show average change over 4 biological replicates, bar shows average of all genes, grey shading indicates less than a 2-fold change in response to dexamethasone. (H-I) Dexamethasone-induced change in expression of genes that encode transcription factors classified as (H) C2H2-type zinc finger proteins or (I) Homeobox. Symbols show the data points from each biological replicate, bars show mean and SEM, grey shading indicates less than a 2-fold change in response to dexamethasone.

We next focused on genes that were differentially expressed in response to dexamethasone compared to a vehicle control (defined as a >2-fold change). We identified 157 differentially expressed genes (DEGs) from cells co-transfected with GR and vector, 99 DEGs from cells co-transfected with GR and FABP1, and 44 DEGs from cells transfected with GR and FABP4. Of these, 84 were unique to GR/vector, 28 were unique to GR/FABP1 and only 5 were unique to GR/FABP4 (Fig 6C). Although only 5 unique genes were identified as DEGs for the GR/FABP4 condition, when we compared all genes in the DEG list across all conditions, hierarchical analysis showed greater similarity between GR/vector and GR/FABP1 transfected cells, compared to GR/FABP4 transfected cells (Fig 6D). There was no change in the classification of the genes across different biological processes when grouped by Gene Ontology terms (Fig S7A).

Interestingly, when examined as a whole, the DEGs that were up-regulated by dexamethasone in GR/vector or GR/FABP1 cells appeared to be unchanged in GR/FABP4 cells (Fig 6E-F). In contrast, overall the DEGs that were upregulated in GR/FABP4 cells were also upregulated in GR/vector and GR/FABP1 cells (Fig 6G). The same trend was observed for DEGs that were down-regulated (Fig S7B-D). This suggests that the effect of dexamethasone may be more limited in cells co-transfected with GR/FABP4 compared to GR/vector or GR/FABP1. We tested this hypothesis by focusing on transcription factors and regulators that were differentially expressed in response to dexamethasone: genes that encode C2H2-type zinc finger proteins (Fig 6H), homeobox proteins (Fig 6I) and predicted transcriptional regulators (Fig S7E). All these genes were up- or down-regulated in response to dexamethasone in control GR/vector cells. In GR/FABP4 cells there was no longer any change in response to dexamethasone in the expression of 8 out of 8 genes encoding C2H2-type zinc finger proteins (Fig 6H), 6 out of 10 genes encoding homeobox proteins (Fig 6I) and 11 out of 15 genes encoding other predicted transcriptional regulators (Fig S7E). Therefore, FABP4 co-expression reversed the effect of dexamethasone for 76% genes identified as transcription factors. In contrast, FABP1 co-expression only reversed the effect of dexamethasone for 36% genes identified as transcription factors.

Together, this suggests that while FABP4 limits the transcriptional activity of the GR for many genes, it enhances the activity of some genes that are up-regulated by classical GRE-controlled transcription.

## Discussion

It is well recognised that FABPs can affect the transcriptional activity of the PPAR-NHR family^11, 12, 13, 14, 15, 16^. PPARs are classified as type II NHRs and their cellular localisation is restricted to the nucleus. Here, for the first time, we show that FABPs can also affect the transcriptional activity of a type I NHR, the GR. Type I NHRs, such as the GR, are generally activated in the cytosol before relocating to the nucleus. We report an increase in GRE-mediated transcriptional activity that occurs in a ligand- and FABP-specific manner. Unlike the PPAR system, where PPAR ligands can cause a conformational change in the FABP to affect NHR activity, we find that FABPs only bind GR ligands at very low affinity (high micromolar to millimolar range). Nevertheless, FABP-dependent increases in GRE-mediated transcription are associated with increased proximity between the GR and FABP in the nucleus (as determined by FRET) and a decrease in the rate of nuclear egress of both proteins (as determined by FRAP). While dexamethasone activation of GR in the presence of FABP caused increased transcription of genes in a GR signalling array, at the level of the transcriptome we instead observed a reversal of the effect of dexamethasone on GR alone. This suggests that differential expression of FABPs can shape the cellular response to activation of the GR.

Both FABPs and GR can shuttle between the nucleus and cytosol^7, 8, 11, 13, 15^. As such, it is the relative rates of import and export from the nucleus that determine where most of the GR or FABP is located within the cell. Addition of ligand appears to stabilise an interaction between GR and FABPs (in a ligand-dependent manner) in the nucleus. This corresponds with a decrease in the nuclear egress of both proteins, and ultimately results in their nuclear accumulation. The nuclear retention of GR has previously been associated with increased gene transcription^24^. As such, the increased retention of the GR in the nucleus in the presence of FABPs could, in part, contribute to the enhanced transcriptional activity we observe. We have previously reported a similar effect of FABPs on the nuclear retention, and therefore transcriptional activity, of PPARα^11^. PPARα is a NHR that is restricted to the nucleus. For this receptor class, FABPs have long been recognised to play a key role in the transport of poorly water-soluble ligands, as well as endogenous fatty acids, to the nuclear-restricted receptors. More recently, FABPs have been found to modulate the transcriptional activity of the NHR in a ligand- and FABP-specific manner. The ability of different FABPs to increase the transcriptional activity of PPARα is also linked to a decreased rate of nuclear egress^11^. However, it has been very difficult to differentiate the lipid-trafficking role of FABPs, from their ability to modulate PPAR transcriptional activity. The discovery that FABPs can also modulate the activity of the GR, with very limited affinity for GR ligands themselves, allows us to differentiate between these two cellular roles for the first time. This separation of FABP activity can also be linked to the different mechanisms that underlie nuclear accumulation of FABP4. Increased PPARγ activity that is dependent on FABP4 is linked to increased nuclear entry^13, 15^, however we find that increased GR activity that is similarly dependent on FABP4 is linked to decreased nuclear egress. Binding of PPARγ ligands to FABP4 causes a change in conformation of residues in the helical cap of FABP4. This rearrangement in the helical cap has been hypothesised to expose a conformational nuclear localisation sequence that increases nuclear entry of the ligand-bound FABP4^13, 15^, inherently linking this mechanism with ligand trafficking. As GR ligands do not bind FABPs with high affinity, this site may remain hidden. As such, decreased nuclear egress of FABP4 could be exclusively linked to a non-trafficking, transcriptional modulation activity of FABPs. However, how FABPs interact with the NHR transcriptional machinery to increase activity, is currently unknown.

DNA sequences can allosterically modify the conformation, and therefore activity, of transcriptional regulators^25^ – analogous to a ligand-receptor binding event dictating activation of intracellular signalling. In the case of the GR and the estrogen receptor (ER), the DNA sequence bound by the NHR can allosterically influence receptor binding partners^26, 27, 28^. Similarly, binding partners of transcriptional regulators will feedback to influence the outcome of DNA binding, to either activate or repress transcription^25^. This is again analogous to intracellular signalling proteins influencing the affinity of receptors for ligand. The allosteric relationship between DNA and NHR seems to underlie the FABP/GR interaction detailed herein. It provides an explanation for the ligand-dependent effects of FABPs on GRE-mediated gene transcription; even though FABPs are unable to directly bind GR ligands, the conformation of the GR induced by ligand binding could dictate whether FABPs can interact with the GR transcriptional machinery. Moreover, it provides an explanation for why FABP4 can promote GRE-mediated transcriptional activity (Fig 1,2,5), but have a repressive effect on GR activity over the whole transcriptome (Fig 7). The GR can affect transcriptional activity by multiple mechanisms. Many genes regulated by the GR have been associated with GRE sites ^10^. However, the GR can also bind to GRE-half sites (as a monomer, rather than a dimer), and influence the activity of other transcription factors^10^. We therefore hypothesise that the conformation of the GR transcriptional machinery induced by FABP4 interaction, could limit the ability of the GR to activate non-GRE-linked target genes.

The GR is expressed ubiquitously throughout the body, which contributes to the wide range of well-documented adverse effects of prolonged glucocorticoid therapy, including osteoporosis, hyperglycaemia, cardiovascular disease, growth retardation, cushingoid appearance (redistribution of fat) and neuropathy^1^. In contrast to the GR, different FABP sub-types have a relatively restricted expression. For example, FABP1 is expressed in the liver, GI tract and kidney and FABP4 is expressed in adipose and breast tissue^29^. The glucocorticoid response genes identified in the liver, hypothalamus and pituitary appear to be most closely associated with GR’s adverse effects^30^. Therefore, it may be that a glucocorticoid that is unable to induce a GR conformation capable of recruiting FABP to the transcriptional machinery, may have a more limited side effect profile. The GR has a long history in the pharmaceutical industry but only modest improvements in this drug class have occurred^30^. The difficulty in fully exploiting NHRs for drug discovery lies in the challenge associated with identifying ligands that only control the desired, subset of therapeutically relevant genes^31^. The ability of FABPs to selectively promote (and in some cases, limit) the transcriptional activity of the GR in a ligand-dependent manner may therefore have implications for drug discovery. Given the increasing evidence for ligand-directed recruitment of co-factors to the GR transcriptional machinery, this property might usefully be included in ligand profiles during drug candidate selection and progression processes.

## Methods

### Plasmids

Human FABP1, FABP2 and GFP-FABP1 have been described previously^11^. The GRE-SEAP reporter gene was from Clontech^32^ and the β-galactosidase reporter vector was from Promega. Human FABP3, human FABP4, human FABP5 and human GR (α sub-type) were synthesized by GeneArt (Germany). All four genes were subcloned into the pSG5 mammalian expression vector purchased from Agilent Technologies. Human FABP4 was subcloned into an expression vector containing green fluorescent protein (GFP), pAcGFP-C1, to make GFP-FABP4. Human GR was subcloned into an expression vector containing dsRed fluorescent protein, pDsRed-Express-N1 (Clontech), to make GR-dsRed.

### FABP expression and purification

FABP1 and FABP2 were expressed and purified as described previously^11, 33^. Codon optimised genes for human FABP3, FABP4 and FABP5 were synthesised by Life Technologies (Thermo Fisher, Australia) and cloned into pET-28a (+) vector between Nco1 and BamH1 sites, to generate N-terminal hexahistidine-tagged proteins with a TEV protease cleavage site. Recombinant FABP3, 4 and 5 were expressed and purified from BL21 (DE3) gold *E.coli* cells as previously described for FABP1 and FABP2^11, 33^, with a few minor modifications. Briefly, the cells were grown in autoinduction media containing kanamycin (50 μg/ml) at 37°C for 3 h. At an optical density (A_600_) of approximately 0.6, the temperature was lowered to 20°C and protein expression induced for approximately 20 h in a shaking incubator (180 rpm). The cells were harvested by centrifugation (5000x g for 20 min). Cells were lysed by sonication in 25 mM HEPES pH 8.0, 250 mM NaCl, 5 mM imidazole, complete EDTA-free protease inhibitor cocktail (Roche) and 10 *μ*g/ml DNAase. Following sonication, cell debris were removed by centrifugation (25000xg for 30 min at 4 °C). FABPs were purified by nickel metal affinity chromatography followed by hydrophobic interaction chromatography. The purified proteins were incubated with TEV protease (at a TEV to FABP ratio of 1:35) at 25 °C for 16 h and the cleaved N-terminal hexahistidine tag was removed using nickel metal affinity chromatography. FABPs were delipidated using a Lipidex-5000 resin (Sigma Aldrich) and buffer exchanged into 20 mM HEPES pH 8.0 and 50 mM NaCl. Protein purity was assessed by SDS-PAGE and correct protein folding was confirmed by ^1^H 1D NMR spectroscopy^12^.

### Fluorescence polarisation binding assays

A fluorescence polarisation binding assay was used to determine the affinity of GR ligands for FABPs. Binding of FABP to a fluorescently-labelled fatty acid, BODIPY 500/510 C4, C9 (Invitrogen), results in reduced tumbling rate of the fluorescent probe and therefore an increase in fluorescence polarisation. Purified and delipidated FABP, 200 nM BODIPY C4, C9, and GR ligand were diluted in 20 mM HEPES pH 8.0, 50 mM NaCl and 2.5 % (v/v) DMSO, in triplicate in 384-well black polystyrene plates (Greiner) with a total assay volume of 36 μL. After equilibration for 1 hr at 37°C, fluorescence polarisation was measured at 37°C using a PHERAstar FS (BMG Labtech) plate reader with fluorescence excitation and emission wavelengths set at 485 nm and 520 nm, respectively. The affinity of FABPs 1-5 for BODIPY C4, C9 were first determined by saturation binding experiments. Increasing concentrations of purified FABPs were added to wells containing 200 nM BODIPY C4, C9 to derive the following K_d_ values: 0.09 μM for FABP1, 4.1 μM for FABP2, 0.02 μM for FABP3, 0.42 μM for FABP4 and 0.16 μM for FABP5. A competition ligand displacement fluorescence polarisation assay was then used to determine the inhibition constants (K_i_) of GR ligands for FABPs. Each well contained 200 nM BODIPY C4,C9, FABPs 1, 2, 3, 4 or 5 at five times the concentration of their respective K_d_ for BODIPY C4, C9, and increasing concentrations of GR ligands (from 10 nM to 1 mM). To obtain the percentage of displacement, control wells containing no compound (zero displacement) and a control compound (maximum displacement) were also included. The control compounds used were: 25 μM GW7647 for FABP1, 500 μM sodium oleate for FABP2, 250 μM WY14643 for FABP3, 250 μM L165-041 for FABP4 and 250 μM GW1929 for FABP5. Fluorescence polarisation was corrected by subtraction of background fluorescence signal and expressed as a percentage of the maximal fluorescence polarisation (%F_Max_) obtained from saturation binding experiments. Competition binding data were analysed using a non-linear regression one-site competitive binding equation in GraphPad Prism (version 7.0) to obtain the K_i_. Data are expressed as mean ± SEM from 3 individual measurements performed in triplicate.

### Radioligand binding assays

Binding to the GR was determined using soluble GR prepared from the cytosol of the IM9 human lymphoblast cell line, as previously described^34^. Competition binding experiments used 1.5 nM [^3^H]-dexamethasone, and increasing concentrations of unlabelled dexamethasone (300 pM – 10 μM), cortisone (3 nM – 100 μM) or methylprednisolone (300 pM – 10 μM). Non-specific binding was determined by 10 μM triamcinolone. Samples were incubated for 6 hours at 4°C and counted using a liquid scintillation counter. Competition binding data were analysed using non-linear regression, with Ki values calculated using the Cheng-Prusoff equation in SigmaPlot 4.0 (SPSS Inc.). Data are expressed as the pKi, with mean ± SD from 2 individual experiments.

### Cell culture and transfection

COS-7 cells were kindly provided by Prof. Phillip Nagley (Monash University, Victoria, Australia). The cells were cultured in Dulbecco’s modified Eagle’s medium with 4 mM glutamine, 100 units/mL penicillin, 100 mg/mL streptomycin, and 10% (v/v) fetal bovine serum in a 95% air, 5% CO_2_ atmosphere at 37 °C. For transfections, cells were seeded into 6-well plates at a density of 2×10^5^ cells/well in culture medium devoid of antibiotics. Transfections were carried out using X-tremeGENE 9 DNA transfection reagent according to the manufacturer’s protocol.

### Immunoblotting

Immunoblotting used primary antibodies recognising FABP1 (Abcam (AB76812; rabbit, 1:300), FABP2 (gift from Dr Satoshi Kaiura, Dainippon Sumi-tomo Pharma Co. Ltd., Osaka, Japan; mouse; 1:400), FABP3 (R&D Systems MAB1678; rabbit, 1:200), FABP4 (Abcam AB92501; rabbit, 1:1000), FABP5 (Abcam AB37267; rabbit, 1:200), GR (Abcam AB183127; rabbit, 1:400) or β-actin (Cell Signaling Technology 3700S; mouse, 1:5000). Primary antibodies were detected using the following fluorescent secondary antibodies: goat anti-mouse 680 (LI-COR 926-68070; 1:10,000), goat anti-rabbit 800 (LI-COR 926-32211; 1:10,000).

Proteins were resolved by SDS–polyacrylamide gel electrophoresis using precast 4-15% Mini-PROTEAN TGX gels (Bio-Rad) and transferred to 0.45-mm LF PVDF (low fluorescence polyvinylidene difluoride) membranes (Bio-Rad) using a Trans-Blot SD Semi-Dry Transfer Cell (for 75 min at 10 V; Bio-Rad). Membranes were blocked for 1 hour at room temperature (5% w/v BSA in PBS with 0.1% v/v Tween 20 [PBS-T]) and incubated with primary antibody overnight at 4°C (diluted in 1% w/v BSA). Membranes were washed, incubated with secondary antibody (diluted in PBS-T) for 1 hour at room temperature, and washed. Immunoreactivity was detected by fluorescence using the Odyssey Classic Infrared Imager (LI-COR Biosciences), with resolution set at 169 μm.

### GRE reporter gene assay

Transactivation assays were carried out using COS-7 cells in six-well plates (2×10^5^ cells/well) co-transfected with 0.3 μg of the GRE reporter (GRE-SEAP), 0.15 μg of β-galactosidase (as a transfection efficiency control), 0.05 μg of either empty vector (pSG5) or GR, and 0.3 μg of either empty vector (pSG5), FABP1, FABP2, FABP3, FABP4 or FABP5. 24 h following transfection, cells were treated with vehicle (0.1% v/v DMSO), or increasing concentrations of dexamethasone, hydrocortisone, prednisolone, cortisone, β-methasone, budesonide, methylprednisolone or prednisone for 24 h. Media samples were taken to detect the secreted alkaline phosphatase reporter (GRE-SEAP) before the cells were lysed and assayed for β-galactosidase activity using the β-galactosidase enzyme assay system (Promega) to correct for differences in transfection efficiency. Media samples were added to 96-well white optiplates (PerkinElmer) in duplicate. Samples were heat-treated (65°C for 30 min) to denature any endogenous alkaline phosphatase and then cooled on ice before addition of SEAP buffer (0.39 M diethanolamine, and 0.98 mM MgCl2, pH 10.3). The fluorescent substrate 4-MUP (0.2 mM) (Sigma-Aldrich) was incubated with the samples for 1 h in the dark. Plates were then read on a Fusion-α microplate reader (PerkinElmer) with excitation at 360 nm and emission at 440 nm. The data are expressed relative to vehicle-treated control, from three independent experiments conducted in duplicate.

### FABP nuclear translocation assay

The Operetta high-content imager (PerkinElmer Life Sciences) equipped with a live cell chamber was used to obtain unbiased images of the localization of GFP-tagged FABPs for automated quantification in live cells. COS-7 cells were co-transfected with 0.05 μg of empty vector (pSG5) or 0.05 μg of GR plasmid DNA and 0.3 μg of GFP-FABP1 or GFP-FABP4 plasmid DNA in six-well plates. Twenty-four hours following transfection, cells were subcultured into black-walled, optically clear 96-well plates and treated for 24 h with vehicle (0.1% v/v DMSO), 10 μM dexamethasone, 1 mM hydrocortisone, 100 μM prednisolone, 1 mM cortisone, 10 μM β-methasone, 10 μM budesonide, 100 μM methylprednisolone or 100 μM prednisone (all at EC_80_ concentrations, defined by GRE reporter gene assay concentration-response curves). On the day of imaging, cells were incubated with Hoechst stain (2 μg/ml) for 5 min at room temperature (to stain the nuclei), prior to imaging the live cells in Hanks’ balanced salt solution at 37 °C. The centre of each well of a 96-well plate was imaged with an Olympus LUCPlanFLN ×20 (NA 0.45) objective, and the nuclear redistribution of FABPs was determined by measuring the fluorescence intensity in the nucleus and cytosol of every cell within the field of view using automated protocols within the Harmony High Content Imaging and Analysis software (version 3.5.2). For each experimental repeat, ~50 cells were analyzed. Experiments were performed on four separate occasions.

### GR-FABP micro-FRET timecourse

COS-7 cells were co-transfected with 0.05 μg of GR or GR-dsRed plasmid DNA and 0.3 μg of FABP1, GFP-FABP1, FABP4 or GFP-FABP4 plasmid DNA in six-well plates. Twenty-four hours following transfection, cells were sub-cultured into black-walled, optically clear 96-well plates and left for 24 h. FRET images were collected using an Operetta high-content imager (PerkinElmer Life Sciences) equipped with a live cell chamber and Olympus LUCPlanFLN ×20 (NA 0.45) objective. Images were acquired and analyzed according to the three-cube micro-FRET method^35, 36^. In brief, a series of live cell images were first collected from cells expressing GFP-FABP and untagged GR or GR-dsRed and untagged FABP to calculate values for GFP bleed-through into the dsRed channel (3.3%) and GFP cross-excitation of dsRed (6.9%) for our imaging system. To calculate FRET in cells co-expressing GR-dsRed and GFP-FABP1 or GFP-FABP4, five separate live cell images were collected: (i) brightfield to define cell region, (ii) GFP using 460-490 excitation filter and 500-550 emission filter, (iii) dsRed using 520-550 excitation filter and 560-630 emission filter, (iv) GFP/dsRed uncorrected FRET using 460-490 excitation filter and 560-630 emission filter, and (v) Hoescht nuclear stain using 360-400 excitation filter and 410-480 emission filter. A baseline image was collected, cells were treated with vehicle (0.1% v/v DMSO), 10 μM dexamethasone, 1 mM hydrocortisone or 100 μM prednisolone, and then images were collected every 1 h for 24 h. For all image sets, 4 fields of view were captured from each well at each time point.

Harmony High Content Imaging and Analysis software (version 4.8) was used to automatically detect analysis regions in each image from each field of view. The brightfield images were used to define the “cell” region, “background” (no cells) was defined by subtracting the cell region from the whole field of view, “nucleus” was defined using the Hoescht nuclear stain images, and “cytosol” was defined by subtracting the nuclear region from the cell region. The fluorescent intensities for GFP, dsRed and GFP/dsRed uncorrected FRET in each region (cytosol, nucleus), and the number of cells analysed (nuclei count using Hoescht nuclear stain images) were automatically calculated by the software. The change in FRET in response to treatment was then calculated. All intensity values were background-subtracted. To correct for GFP bleed-through and cross-excitation of dsRed, 3.3% of the GFP intensity and 6.9% of the dsRed intensity were subtracted from the GFP/dsRed FRET intensity to produce a corrected pseudocolor FRET image, FRETc. To quantify the degree of sensitized FRET observed between GFP-FABP1 and GR-dsRed compared to GFP-FABP4 and GR-dsRed, average intensities of the FRETc images were divided by the product of the GFP and dsRed image intensities to produce normalized NFRETc values^35, 36^. For each experimental repeat at least 200 cells per condition were analysed from 4 fields of view. Experiments were performed on 3 separate occasions. Representative pseudocolour corrected FRET images were generated by using the “FRET and Colocalisation Analyser” plugin^37^ for the Fiji distribution of Image J^38^, using image regions with good co-localisation of GFP and dsRed fluorescence. For each condition, all four fields of view were used to calculate the mean donor and acceptor bleedthrough using the plugin.

### Fluorescence recovery after photobleaching (FRAP)

COS-7 cells in 6-well plates were co-transfected with 0.3 μg/well of FABP1, GFP-FABP1, FABP4 or GFP-FABP4 and 0.05 μg/well of GR or GR-dsRed plasmid DNA. Twenty-four hours following transfection, cells were sub-cultured into Ibidi μ-Slide 8-well chambered coverslips and left for 24 h. On the day of the experiment, cells were treated with vehicle (0.1% v/v DMSO) or 10 μM dexamethasone for 6 h at 37 °C prior to imaging on a Leica Sp8 confocal microscope with a HCX PL APO 63x (NA 1.40) oil objective. FRAP experiments were performed and data analysed using the FRAP wizard in Leica LAS AF software. FRAP was performed by either bleaching the nucleus and monitoring fluorescence recovery back into the bleached region to determine nuclear entry, or by bleaching the cytosol and monitoring fluorescence recovery back into the bleached region to determine nuclear egress. Fluorescence recovery curves (F_Rec_), maximal fluorescence recovery (FRAP Max) and the time to half fluorescence recovery (FRAP t_1/2_) were all calculated using the LAS AF FRAP wizard. For each experimental repeat 6 cells were analysed. Experiments were performed on 3 separate occasions.

### qPCR array and analysis

COS-7 cells were co-transfected with 0.1 μg of GR plasmid DNA and 0.6 μg of either empty vector (pSG5), FABP1 or FABP4 plasmid DNA in 6-well plates. Twenty-four hours following transfection, cells were treated with vehicle (0.1% v/v DMSO) or 10 μM dexamethasone for 24 h. RNA was extracted using the RNeasy mini kit (Qiagen), cDNA was generated using the RT2 First Strand Kit (Qiagen). qPCR was performed using the RT^2^ SYBR Green Mastermix and Glucocorticoid Receptor Signalling RT^2^ Profiler PCR Array from Qiagen, according to the manufacturer’s instructions, and a Bio-Rad CFX96 Touch Real-Time PCR Detection System. The change in Ct value relative to vehicle control and housekeeping genes (HPRT1 and RPLP0; ΔΔCt) was calculated. Changes at least 2-fold compared to vehicle control were considered significant. Venn diagrams were created using the custom Venn diagram tool by Bioinformatics and Evolutionary Genomics (http://bioinformatics.psb.ugent.be/webtools/Venn/). Experiments were performed on 3 separate occasions.

### RNA sequencing and analysis

COS-7 cells were co-transfected with 0.1 μg of GR plasmid DNA and 0.6 μg of either empty vector (pSG5), FABP1 or FABP4 plasmid DNA in 6-well plates. Twenty-four hours following transfection, cells were treated with vehicle (0.1% v/v DMSO) or 10 μM dexamethasone for 24 h, RNA was extracted using the RNeasy mini kit (Qiagen). Cell transfection, treatment and RNA extraction was performed on four separate occasions. Transcriptome sequencing and bioinformatic analysis was performed by the Beijing Genomics Institute (BGI).

All data were expressed relative to vehicle-treated controls, ratios were log2 normalized to allow quantitative analysis, and any nonvalid values were removed. Genes with at least a 2-fold change compared to vehicle control were classified as differentially expressed genes (DEGs) and were retained for further analysis. DEGs were identified by BGI bioinformatic analysis using DESeq2. Ingenuity Pathway Analysis (IPA; Qiagen, Summer 2020 release) was used to determine the most likely upstream regulator of the DEGs in the GR/pcDNA control group. Hierarchical clustering of DEGs was performed using IPA (default settings). Venn diagrams were created using the custom Venn diagram tool by Bioinformatics and Evolutionary Genomics (http://bioinformatics.psb.ugent.be/webtools/Venn/). BGI bioinformatic analysis identified genes that were predicted to encode transcription factors and classified DEGs according to their biological process Gene Ontology (GO) term. To generate Biological Process GO pie charts, classifications with P < 0.05 were used to group proteins according to biological function, synonymous classifications were removed, and the number of proteins classified within these groups was counted.

## Acknowledgements

These studies were supported by an Australian Research Council (ARC) grant DP150102587. Michelle Halls was supported by a National Health and Medical Research Council of Australia (NHMRC) RD Wright Fellowship 1061687, a Monash Strategic Bridging Fellowship, and a Viertel Senior Medical Research Fellowship (from The Cross Family and Frank Alexander Trusts).

