## Supplementary Information for "Fatty acid binding proteins shape the cellular response to activation of the glucocorticoid receptor"

**Supplementary Table.**

**Table S1. Binding affinity of GR ligands (endogenous and synthetic) for GR.** Competition binding of GR ligands at the glucocorticoid receptor (GR). The table shows pKi values as mean and standard deviation (n=2).

| Ligand | GR |
|---|---|
| Dexamethasone | 8.32±0.04 |
| Cortisone | 5.84±0.04 |
| Methylprednisolone | 8.56±0.05 |

### Supplementary Figures.

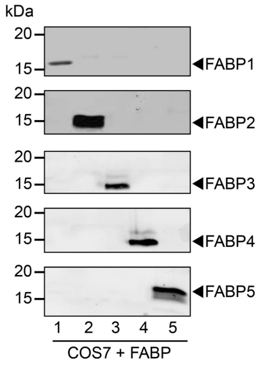


**Figure S1. COS-7 cells do not endogenously express FABPs.** Expression of FABP1-5 detected by immunoblotting following transfection of COS-7 cells.

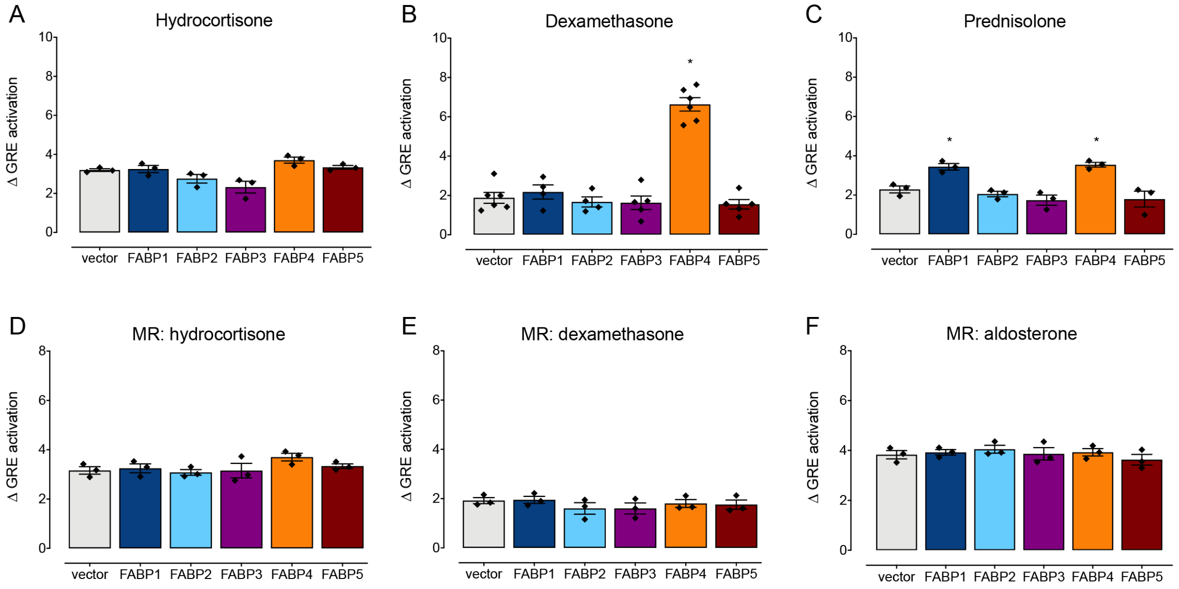


**Figure S2. GRE activity is not affected by the MR in COS-7 cells.** GRE transcription in COS-7 cells transfected with GR and vector control, FABP1, FABP2, FABP3, FABP4 or FABP5 in response to an $EC_{80}$ concentration of (A) hydrocortisone, (B) dexamethasone or (C) prednisolone (n=3-6). GRE transcription in COS-7 cells transfected with MR and vector control, FABP1, FABP2, FABP3, FABP4 or FABP5 in response to (D) hydrocortisone, (E) dexamethasone or (F) aldosterone (n=3). Bars show mean, error bars SEM and data points show individual experiments. * $p<0.05$ versus vector control, one-way ANOVA with Dunnett's multiple comparisons test.

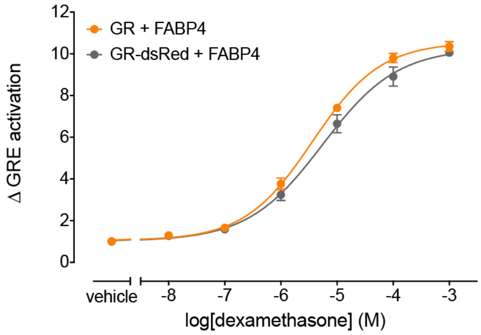


**Figure S3. GR-dsRed validation.** GRE reporter gene concentration-response curves to dexamethasone for untagged GR and GR-dsRed (n=3). Symbols are means and error bars SEM.

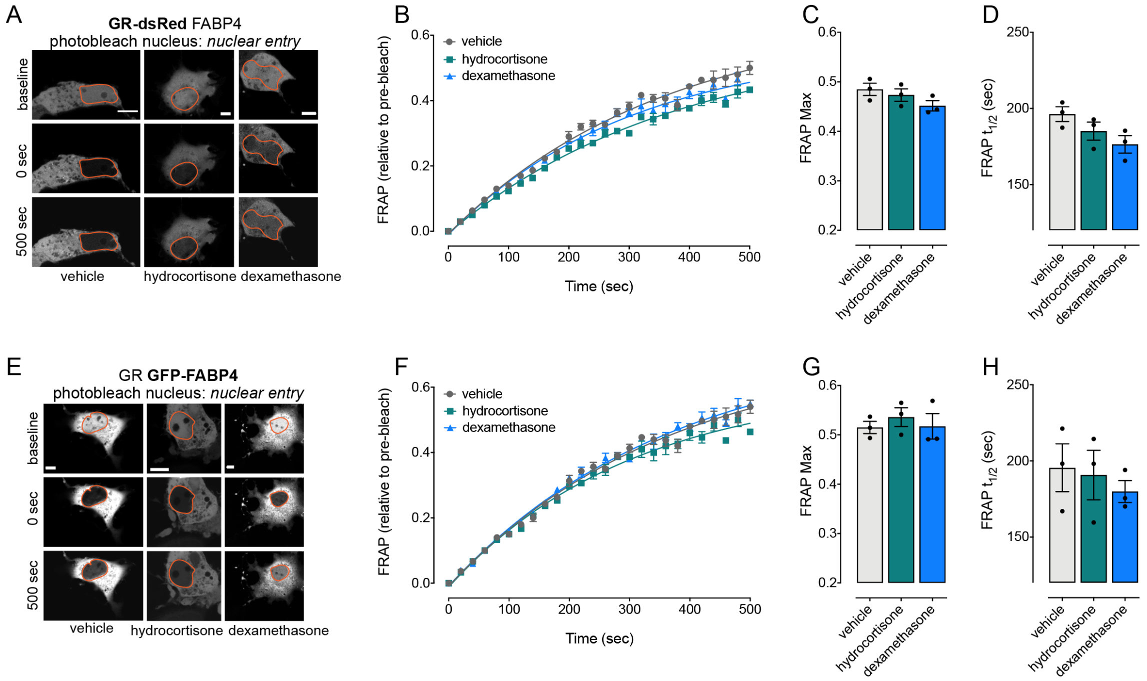


**Figure S4. No change in nuclear entry.** COS-7 cells were treated with vehicle, hydrocortisone or dexamethasone for 6 h, before photobleaching of the nucleus. (A-D) GR mobility was monitored in cells co-transfected with GR-dsRed and FABP4 (n=3). (E-H) FABP4 mobility was monitored in cells co-transfected with GR and FABP4-GFP (n=3). A,E show representative images with the nucleus outlined in orange. Scale bar is 10 μm. B,F show FRAP time course, symbols are means and error bars SEM. C,D show maximum recovery. G,H show the half time of recovery. Bars are means, error bars SEM and symbols show individual data points for each experiment.

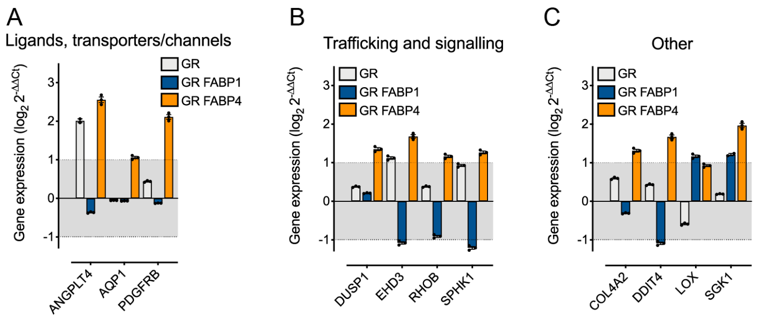


**Figure S5. FABP4 increases GR-mediated gene transcription in response to dexamethasone.** Genes with an average 2-fold change in expression in cells transfected with GR and FABP4 in at least 2 experiments, classified according to function. (A) Genes that encode ligands, transporters and channels. (B) Genes that encode proteins involved in intracellular trafficking and signalling. (C) Genes that encode proteins involved in the stress response, and regulation of extracellular collagen. Bars are means, error bars SEM and symbols show individual data points for each experiment. Shaded area (between -1 and 1) indicates genes with less than a 2-fold change in response to dexamethasone.

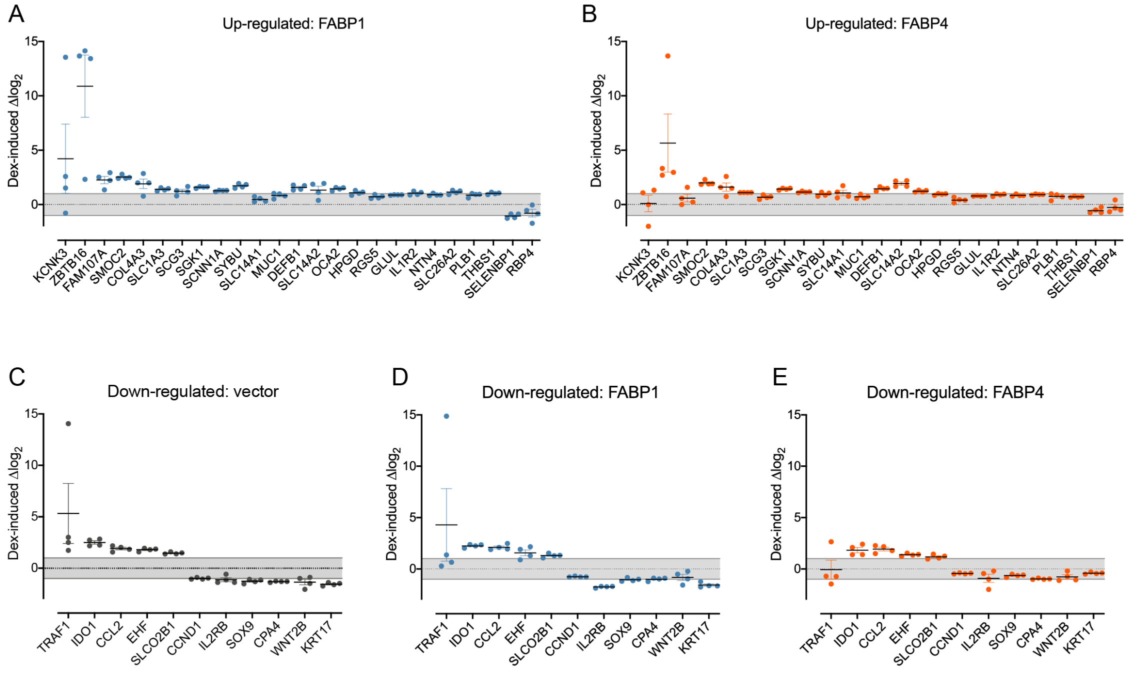


**Figure S6. Validation of transcriptomic dataset.** The dexamethasone-induced change in genes that were identified as dexamethasone-signature genes by Ingenuity Pathway Analysis (IPA) software (n=4). (A-B) Dexamethasone-induced change in expression of genes that were expected to be up-regulated by dexamethasone in COS-7 cells co-transfected with (A) GR/FABP1 or (B) GR/FABP4. (C-E) Dexamethasone-induced change in expression of genes that were expected to be down-regulated by dexamethasone in COS-7 cells co-transfected with (C) GR/vector, (D) GR/FABP1 or (E) GR/FABP4). Symbols show the data points from each biological replicate, bars show mean and SEM, grey shading indicates less than a 2-fold change in response to dexamethasone.

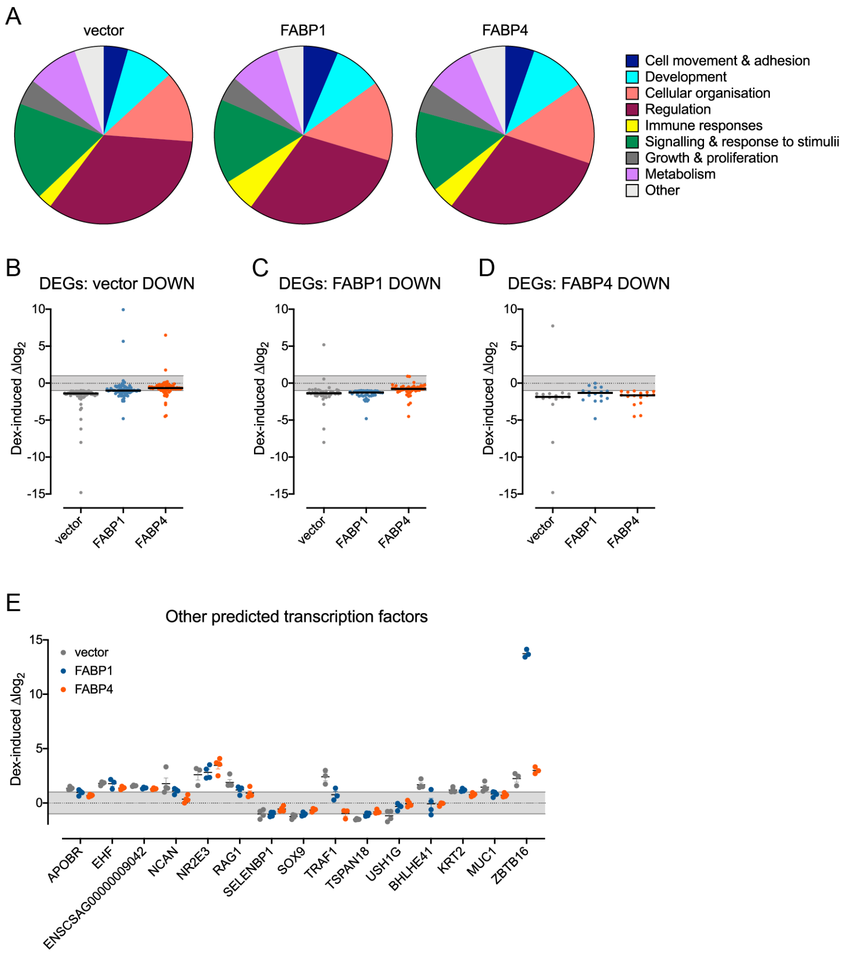


**Figure S7. Altered transcription profile of dexamethasone-stimulated GR in the presence of FABP4.** (A) Genes with a >2-fold change in response to dexamethasone in cells co-transfected with GR/vector, GR/FABP1 or GR FABP4 were classified according to their Gene Ontology biological process term using DAVID. (B-D) The average change in differentially expressed genes (DEGs) in response to dexamethasone across all transfection conditions, with a focus on genes that were down-regulated more than 2-fold in (B) GR/vector cells, (C) GR/FABP1 cells and (D) GR/FABP4 cells. Symbols show average change over 4 biological replicates, bar shows average of all genes, grey shading indicates less than a 2-fold change in response to dexamethasone. (E) Dexamethasone-induced change in expression of genes that encode transcription factors. Symbols show the data points from each biological replicate, bars show mean and SEM, grey shading indicates less than a 2-fold change in response to dexamethasone.
